# Identification of putative reader proteins of 5-methylcytosine and its derivatives in *Caenorhabditis elegans* RNA

**DOI:** 10.1101/2022.01.24.477592

**Authors:** Isabela Cunha Navarro, Kin Man Suen, Dalila Bensaddek, Arun Tanpure, Angus Lamond, Shankar Balasubramanian, Eric A. Miska

## Abstract

Methylation of carbon-5 of cytosines (m^5^C) is a post-transcriptional nucleotide modification of RNA with widespread distribution across organisms. m^5^C has been shown to participate in mRNA transport and maintain mRNA stability through its recognition by the reader proteins ALYREF and YBX1, respectively. We recently showed that m^5^C is required for *Caenorhabditis elegans* development and fertility under heat stress. To contribute to the understanding of how m^5^C and its oxidative derivatives mediate their functions, we developed RNA baits bearing modified cytosines in diverse structural contexts to pulldown potential readers in *C. elegans*. Our mass spectrometry analyses reveal unique binding proteins for each of the modifications. We validate our dataset by demonstrating that the nematode ALYREF homologues ALY-1 and ALY-2 preferentially bind m^5^C *in vitro*. The dataset presented here serves as an important scientific resource that will support the discovery of new functions of m^5^C and its derivatives.

## INTRODUCTION

Nucleoside chemical modifications are a common feature in RNA molecules. More than 160 post-transcriptional modifications have been reported since the discovery of pseudouridine in 1957 (Davis & Allen, 1957; Cohn, 1960; Boccaletto *et al*, 2017). However, the molecular and physiological functions of most RNA modifications remain unknown.

Discovered in 1958 (Amos & Korn, 1958), the methylation of carbon-5 of cytosines in RNAs (m^5^C) is now known to be a conserved and widely distributed feature in biological systems. m^5^C is catalysed by tRNA aspartic acid MTase 1 (DNMT2) and RNA methyltransferases from the Nop2/Sun domain family (NSUN1-7 in humans), and has been shown to be involved in tRNA stability, ribosome fidelity and translation efficiency (reviewed in García-Vílchez *et al*, 2019). Previous studies showed that m^5^C is subject to hydroxylation and oxidation by alpha-ketoglutarate-dependent dioxygenase ABH (ALKBH1), forming 5-hydroxymethylcytidine (hm^5^C) and 5-formylcytidine (f^5^C) (Huber *et al*, 2015; Kawarada *et al*, 2017). Further processing can occur through 2’-O methylation by FtsJ RNA Methyltransferase Homolog 1 (FTSJ1), yielding 2’-O-methyl-5-hydroxymethylcytidine (hm^5^Cm) and 2’-O-methyl-5-formylcytidine (f^5^Cm) (Païs de Barros *et al*, 1996; Huber *et al*, 2017; Kawarada *et al*, 2017). While the biogenesis of these chemical marks has been elucidated (Navarro *et al*, 2020; Kawarada *et al*, 2017), their functions remain largely unexplored.

The identification of readers, *i*.*e*. proteins that specifically, or preferentially, interact with RNA molecules bearing a certain modification, has proven to be an important step towards the understanding of a modification’s biological function. For example, the discovery of N^6^-methyladenosine (m^6^A) readers added new layers of complexity to this pathway, revealing context- and stimuli-dependent functions previously not appreciated (Shi *et al*, 2019). To date, only two reader proteins have been identified for m^5^C. The Aly/REF export factor (ALYREF) was identified in an RNA pulldown experiment and shown to interact with NSUN2-modified CG-rich regions and regions immediately downstream of translation initiation sites. NSUN2-mediated mRNA methylation is interpreted by binding of ALYREF to modulate nuclear-cytoplasm shuttling of transcripts. (Yang *et al*, 2017). More recently, the Y-box binding protein 1 (YBX1) was shown to recognise m^5^C in mRNAs and enhance its stability through the recruitment of ELAV1 in human urothelian carcinoma cells (Chen *et al*, 2019). In zebrafish, YBX1 was shown to stabilise m^5^C-modified transcripts via Pabpc1a during the maternal-to-zygotic transition (Yang *et al*, 2019).

Recently, we used *Caenorhabditis elegans* as a model to engineer the first organism completely devoid of m^5^C in RNA. We mapped m^5^C with single nucleotide resolution in different RNA species and found that m^5^C is required for physiological adaptation and translation efficiency at high temperatures (Navarro *et al*, 2020). Here, to further expand our knowledge on the m^5^C pathway in this organism, we produced the first list of putative readers of m^5^C and its metabolic derivatives hm^5^C, hm^5^Cm and f^5^C. The putative readers are enriched in proteins with roles in germline development and RNA surveillance. We show that the *C. elegans* ALYREF homologues ALY-1 and ALY-2 preferentially bind m^5^C *in vitro* and identify other proteins that contain a conserved RNA recognition motif potentially involved in m^5^C binding.

## RESULTS

To increase the chances to identify *bona-fide* modification readers, we used modified RNA baits with biologically relevant characteristics. Considering that m^5^C is deposited in tRNAs in a structure-dependent manner, we designed baits bearing modifications in three structural contexts: single-strand, double-strand, and loop (**Figure 1A**). m^5^C, hm^5^C, hm^5^Cm and f^5^C monomers were synthesised and incorporated into oligonucleotides tethered to triethylene glycol (TEG)-biotin, as previously described (Tanpure & Balasubramanian, 2017).

**Figure 1.**
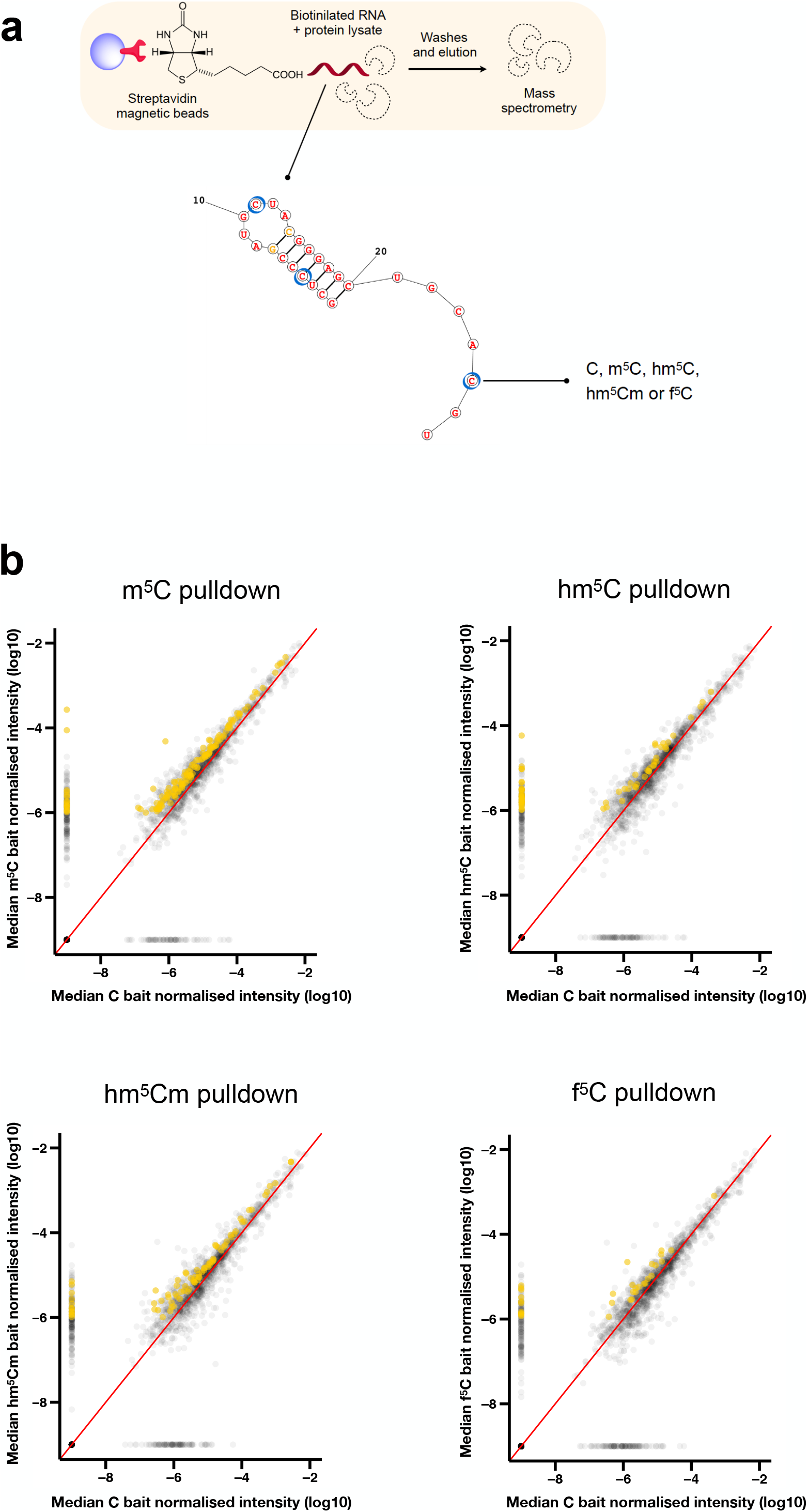
Identification of RNA modification candidate readers. (a) Molecular baits were designed bearing modifications in three structural contexts: single-strand, double-strand, and loop. m^5^C, hm^5^C, hm^5^Cm and f^5^C monomers were synthesised and incorporated into oligonucleotides tethered to TEG-biotin. Blue circles indicate modified positions. Pre-cleared protein lysate from synchronised adult populations was incubated with unmodified or modified biotinilated RNA baits. Streptavidin-conjugated magnetic beads were used to capture proteins that interact with the RNA. Following washes and elution, proteins were processed and identified by RP LC-MS. (b) Scatter plot of proteins bound to modified versus unmodified RNA baits. Median of the normalised iBAQ values were plotted. Proteins uniquely enriched in m^5^C, hm^5^C, hm^5^Cm and f^5^C pulldowns are highlighted in mustard. Proteins were considered candidate readers whenever the fold-change of iBAQ values over the “beads only” and “unmodified C” controls was >2.0 and >1.5, respectively. n = 3 independent biological replicates.

The RNA pulldown experiments were performed in independent biological triplicate. We incubated the RNA baits with pre-cleared whole worm lysates prepared from synchronised populations of gravid adults. To increase the stringency of our screen, we included two controls: beads only and an unmodified RNA bait. Following the pulldown experiments, proteins were eluted from the RNA baits and identified by mass spectrometry.

Proteins were ranked according to their enrichment in modified RNA pulldowns in comparison to the controls. Proteins were considered candidate readers if they preferentially bound to the modified-RNA bait, i.e., at least 2.0 and 1.5 fold increased as compared with the beads and unmodified C controls, respectively (**Figure 1B**).

To narrow down the number of candidate readers, we focused on unique binders, *i*.*e*. proteins that bound specifically to a single modified RNA bait. Using these criteria, we found 143, 62, 81 and 33 putative unique readers for m^5^C, hm^5^C, hm^5^Cm and f^5^C, respectively (**Figure 2A, Supplemental Table S1**). We used the WormBase Enrichment Suite to perform phenotype enrichment analysis on the lists of unique candidates (Angeles-Albores *et al*, 2016). This tool analyses a list of genes according to phenotypes that have been reported by researchers to WormBase upon knockout or RNAi, thus suggesting a potential involvement of such genes in specific biological processes. m^5^C unique binders are the most abundant proteins when compared to its oxidised derivatives (Huber *et al*, 2015, 2017), likely due to higher levels of this modification in RNA. This subset is enriched mostly for germline development phenotypes, and a few RNA surveillance processes. hm^5^Cm-unique binders show specific enrichment for RNA processing phenotypes, whereas hm^5^C and f^5^C show exclusively germline-related phenotypes (**Figure 2B**).

**Figure 2.**
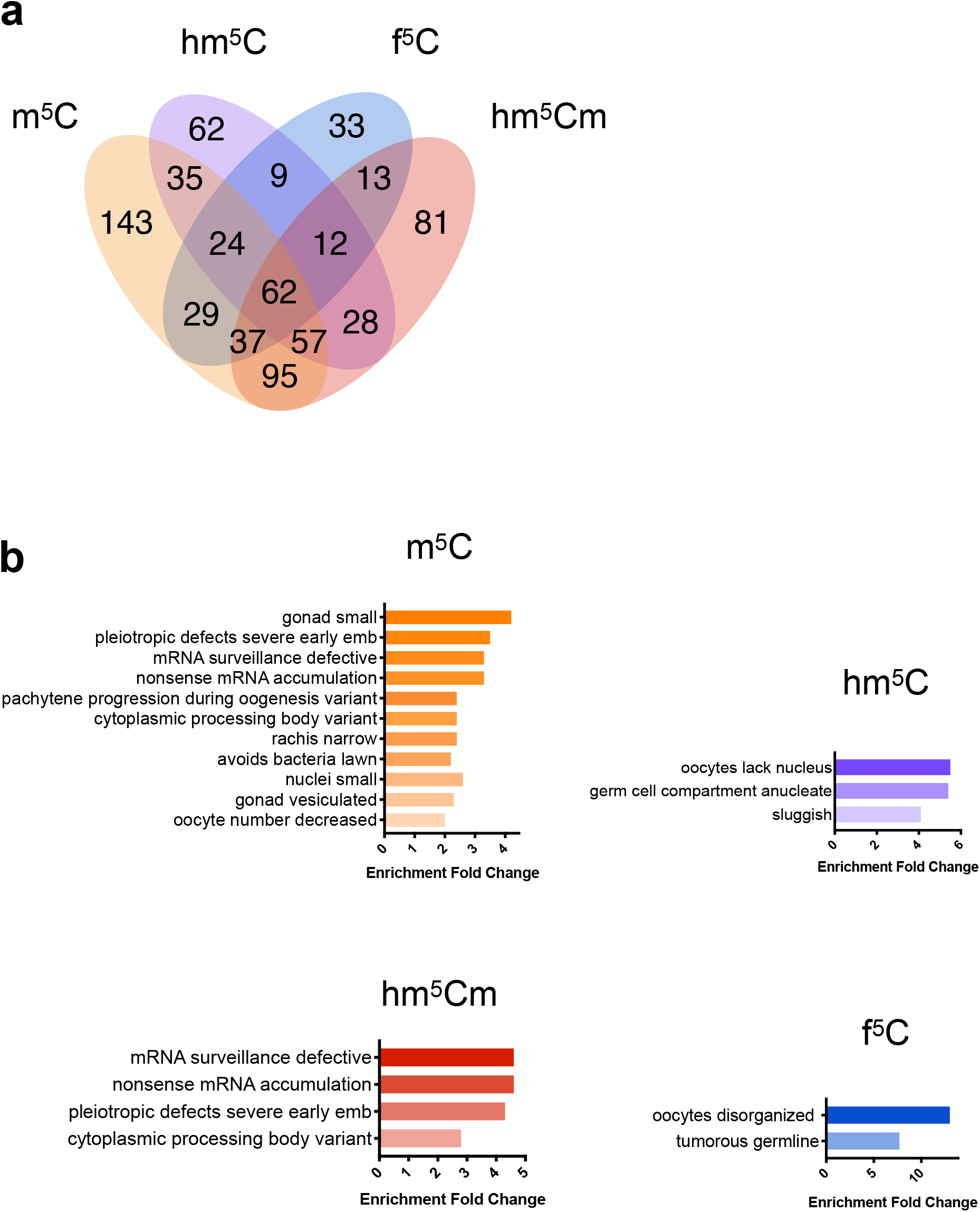
Phenotype enrichment for proteins identified as unique candidate readers. (a) Venn diagram showing the number of enriched proteins detected by mass spectrometry in pulldowns using RNA with different m^5^C, hm^5^C, hm^5^Cm or f^5^C. (b) Phenotype enrichment analysis of unique binders of m^5^C, hm^5^C, hm^5^Cm or f^5^C. Enrichment analysis was performed with the Wormbase Enrichment Suite. Colour code: m^5^C in orange, hm^5^C in purple, hm^5^Cm in red and f^5^C in blue.

To validate our dataset, we verified whether ALYREF homologues could be found among our ranked m^5^C reader candidates. Yang *et al*. identified amino acid residues on ALYREF required for binding to m^5^C (Yang *et al*, 2019). Protein alignments show that some of these residues are conserved in ALY-1 and ALY-2 (**Figure 3A**). Indeed, ALY-1 and ALY-2 are 1.8-fold enriched across all triplicates of m^5^C pulldowns (**Supplemental Table S1**). We used transgenic strains expressing either ALY-1 (OP502), or ALY-2 (OP217), tagged at the C-terminus with TY1::eGFP::3xFLAG, to confirm these findings (Sarov *et al*, 2012). We performed pulldown experiments under the same conditions as before and confirmed using western blotting that both ALY-1 and ALY-2 preferentially bind m^5^C *in vitro* (**Figure 3B**).

**Figure 3.**
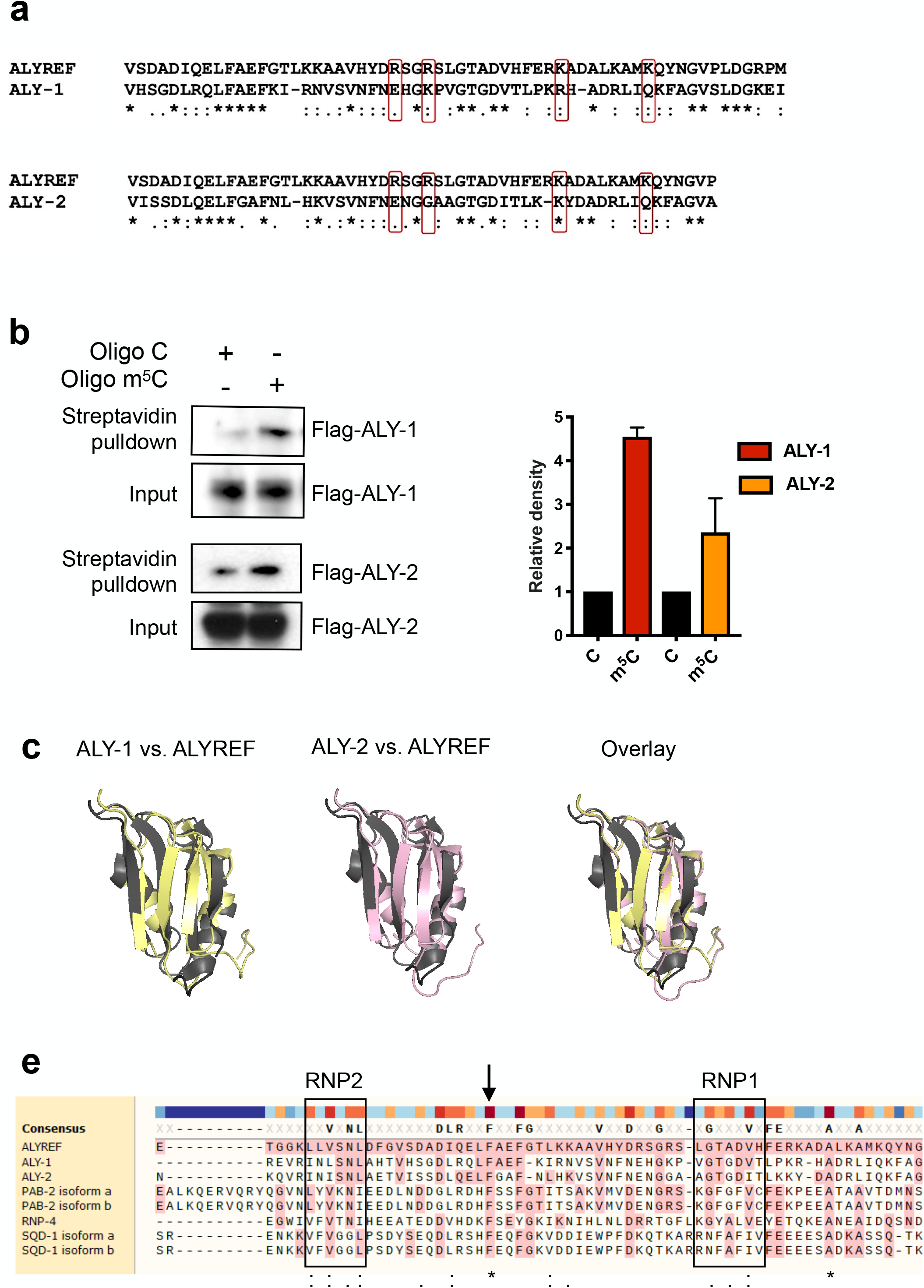
ALY-1 and ALY-2 preferentially bind to m^5^C-modified RNA *in vitro*. (a) Protein sequence alignment of ALYREF and ALY-1/ALY-2 using Clustal Omega. Residues in ALYREF required for m^5^C-binding previously identified by Yang *et al*. are highlighted in red box. “*” represents identical amino acids while “:” and “.” represent amino acids with strongly and weakly similar properties, respectively. (b) Anti-FLAG western blotting (left) showing enrichment of ALY-1 and ALY-2 following pulldown using m^5^C-modified RNA oligos and relative quantification (right). Mean ± SD, quantification performed using ImageJ, n = 2 independent biological replicates. (c) Overlay of ALY-1 (yellow) or ALY-2 (pink) with ALYREF (grey) structures. ALY-1 and -2 structures were predicted using the SWISS-Model platform which were aligned against a ALYREF crystal structure (PDB: 3ULH). The overall RRM folds are conserved in ALY-1 and -2. (d) Sequence alignment of RRM in m^5^C unique binders with ALYREF using Muscle. Black squares highlight consensus sequences RNP1 and RNP2 that are found in RRM. “*” represents identical amino acids while “:” and “.” represent amino acids with strongly and weakly similar properties, respectively. Amino acids highlighted in pink are identical to the reference sequence (ALYREF).

Some m^6^A readers use common RNA binding domains to identify modified RNA (Shi *et al*, 2019). Amino acid residues shown to be important for m^5^C binding in ALYREF also reside in the RNA recognition motif (RMM) domain, which is a known RNA-binding domain (Yang *et al*, 2017). The *C. elegans* orthologues ALY-1 and -2 are predicted to have a similar protein fold in the same region (**Figure 3C**).

Having shown that ALY-1 and ALY-2 preferentially bind m^5^C-modified RNA, we wondered if other RRM-containing proteins were also identified as m^5^C unique binders. We found that three other proteins with one or more RRMs, RNP-4, SQD-1 and PAB-2, bind to m^5^C baits preferentially. RRM domains consist of a four-stranded antiparallel ß-sheet with two α-helices packed against it (Maris *et al*, 2005). Sequence alignments of the RRM domains in ALY-1, ALY-2, RNP-4, SQD-1, PAB-2 and ALYREF show that a high degree of conservation exists across these proteins in the two ribonucleoprotein (RNP) consensus sequences that are important for nucleic acid binding. Interestingly, a phenylalanine residue outside of the two RNPs is highly conserved in all six proteins (**Figure 3D**).

In summary, we have identified putative readers for m^5^C, hm^5^C, hm^5^Cm and f^5^C in *C. elegans* using proteomics. To verify the relevance of this proteomic dataset, we confirmed that ALY-1 and ALY-2 interact with m^5^C orthogonally. Finally, we identified three candidate proteins that share a conserved RRM domain with ALY-1/2 (RNP-4, SQD-1 and PAB-2), highlighting these as potential *bona fide* m^5^C readers.

## DISCUSSION

This work presents a list of candidate readers of the m^5^C oxidative derivatives hm^5^C, hm^5^Cm and f^5^C. Stable isotope labelling experiments suggest that hm^5^Cm is a stable modification (Huber *et al*, 2017). In addition, it has been shown that the ratios between hm^5^C, hm^5^Cm and f^5^Cm in the tRNA Leu-CAA pool varies between organisms and tissues, suggesting a dynamic regulation of these marks in different physiological contexts (Kawarada *et al*, 2017). This could suggest that these modifications carry out cellular functions themselves, rather than being metabolic by-products of a demethylation pathway. Considering the reported function of f^5^C in mitochondrial protein synthesis (Takemoto *et al*, 2009), it is expected that the oxidation of m^5^C in tRNAs might affect their role in cytoplasmic translation. The identification of readers provides a first step to elucidate these functions. Importantly, pathogenic mutations have been mapped to NSUN2, NSUN3 and FTSJ1, suggesting that imbalances in these modifications might contribute to the pathogenesis of human diseases (Abbasi-Moheb *et al*, 2012; Khan *et al*, 2012; Komara *et al*, 2015; Van Haute *et al*, 2016; Guy *et al*, 2015; Takano *et al*, 2008; Ramser *et al*, 2004; Freude *et al*, 2004).

The use of synthetic modified RNAs as a bait for identification of proteins that are either specifically, or preferentially, interacting with ribonucleoside modifications has been applied with success in the past (Dominissini *et al*, 2012; Yang *et al*, 2017; Chen *et al*, 2019; Yang *et al*, 2019). Rather than focusing on sequence context, we developed baits bearing modifications in diverse structural contexts. In this work, we identified two proteins showing higher binding affinity to modified RNA: ALY-1 and ALY-2.

The genome of *C. elegans* encodes three members of the Aly/REF family. It has been shown that simultaneous downregulation of these three genes by RNAi does not compromise either viability, or development. In addition, no defects in mRNA export were observed upon simultaneous knockdown of ALY-1, ALY-2 and ALY-3, suggesting that these proteins mediate alternative processes in the nematode, or that their role in transcript export is redundant with other genes (Longman *et al*, 2003). This is in agreement with reports showing that in *Drosophila* ALY proteins are not required for general mRNA transport (Gatfield & Izaurralde, 2002). In contrast, ALY-1 and ALY-2 have been implicated in nuclear retention of a specific mRNA. In *C. elegans*, sex is initially specified by the ratio of X chromosomes to autosomal chromosomes, *i*.*e*. XX animals are hermaphrodites and X0 are males. Female fate requires the genes *tra-1* and *tra-2*, while male cell fate requires inhibition of *tra-2* activity. Kuersten et al. showed that binding of NXF-2 and ALY-1/2 inhibits nuclear export of *tra-2*. In support of these findings, RNAi against ALY proteins led to 5-7% female occurrence in the progeny (Kuersten *et al*, 2004). It remains to be determined whether m^5^C acts as an intermediate in this process, for example by enhancing binding affinity of ALY proteins to *tra-2* mRNA. Notably, our previous report demonstrated that m^5^C is absent or occurs very rarely in the coding transcripts in *C. elegans*, suggesting that ALY-1/2 could be recognising m^5^C in other molecules (Navarro *et al*, 2020). In addition, the existence of multiple orthologs of ALY genes in *C. elegans* raises the possibility that these proteins could act redundantly or bind differential sets of modified residues in RNA.

Our study presents the first comprehensive investigation of putative readers of m^5^C and its derivatives in a whole organism. This dataset represents an important resource for the discovery of new functions of the RNA m^5^C pathway.

## MATERIALS AND METHODS

### Genetics

*C. elegans* strains were grown and maintained as described in Brenner (Brenner, 1974). The strains were kept at 20°C, unless otherwise indicated. HB101 strain *Escherichia coli* was used as food source (*Caenorhabditis* Genetics Center, University of Minnesota, Twin Cities, MN, USA). Bristol N2 was used as the wild type strain.

### Oligonucleotide synthesis

The synthesis of 2’-O-methyl-5-hydroxymethylcytidine (hm^5^Cm), 5-hydroxymethylcytidine (hm^5^C) and 5-formylcytidine (f^5^C) phosphoramidite monomers was performed as described in Tanpure & Balasubramanian, 2017. RNA oligonucleotides (5’-GCXUCCGAUGXUACGGAGGCUGAXC-biotin-3’, where X = C, m^5^C, hm^5^C, f^5^C or hm^5^Cm) were synthesised in collaboration with ATDBio, Southampton, UK. Purity and integrity of all modified RNA oligonucleotides were confirmed by LC-MS analysis.

### Protein extraction

Nematodes were grown until gravid adults in 140 mm NGM plates seeded with concentrated HB101 *E. coli* and then harvested and washed twice in M9 buffer. A final wash was performed in lysis buffer (20 mM HEPES pH 7.5, 150 mM KCl, 1.5 mM MgCl_2_, 0.1% IGEPAL, 0.5 mM DTT) and the animals were resuspended in 4 ml of lysis buffer before the addition of a protease inhibitor cocktail (SIGMAFAST protease inhibitor tablets, Sigma). The animals were pelleted and as much liquid as possible was removed before the samples were frozen as droplets in liquid nitrogen, using a Pasteur pipette. Frozen droplets were transferred to metallic capsules and cryogenically grinded for 25 sec in a mixer (Retsch MM 400 Mixer Mill). The resulting frozen powder was stored at -80°C until the moment of use. The powder was defrosted at 4°C and the lysate was sonicated for 10 cycles of 20 sec, with breaks of 20 sec in between, in a Bioruptor Pico (Diagenode). The sample was centrifuged at maximum speed for 15 min at 4°C and the protein concentration of the supernatant was determined using a BioRad Bradford protein assay (DTT tolerant).

### RNA pulldown

RNA pulldown protocol was adapted from Dominissini *et al*, 2012. Magnetic beads (Dynabeads MyOne Streptavidin C1, Invitrogen) were washed twice in 100 mM Tris-HCl pH 7.5, 10 mM EDTA, 1 M NaCl, and 0.1% Tween-20, followed by two washes in lysis buffer. In order to reduce non-specific protein binding to the RNA bait matrix, lysates were pre-cleared by incubation with washed beads for 1 h at 4°C with constant rotation prior to the pulldown. Finally, the following pulldown mix was prepared: 2 mg of protein lysate, 50 U of RNAse inhibitor Superase-IN (Invitrogen), 500 pmol of 25-mer biotinylated RNA bait, up to a final volume of 250 μl in lysis buffer. The mixture was incubated with constant rotation for 2 h at 4°C and then added to 20 μl of streptavidin-conjugated beads for pulldown for another 2 h at 4°C. The beads were washed 3-5 times in 1 ml of lysis buffer and the bound proteins were eluted for mass spectrometry in 50 μl 4% SDS 100 mM TEAB at 95°C for 10 min.

### Sample preparation for protein mass spectrometry

Samples were processed and analysed as described in Bensaddek *et al*, 2016. Samples were reduced using 25 mM tris-carboxyethylphosphine TCEP for 30 min at room temperature, then alkylated in the dark for 30 min using 50 mM iodoacetamide. Protein concentration was quantified using the EZQ assay (Thermo Fisher). The lysates were diluted with 100 mM triethyl ammonium bicarbonate (TEAB) 4-fold for the first digestion with endoprotease Lys-C (Fujifilm Wako Chemicals), then further diluted 2.5-fold before a second digestion with trypsin. Lys-C and trypsin were used at an enzyme to substrate ratio of 1:50 (w/w). The digestions were carried out overnight at 37°C, then then the tryptic digestion was terminated by acidification with trifluoroacetic acid (TFA) to a final concentration of 1% (v:v). Peptides were desalted using C18 Sep-Pak cartridges (Waters) following manufacturer’s instructions, dried and dissolved in 5% formic acid (FA).

### Reverse-phase liquid chromatography-MS

RP-LC was performed using a Dionex RSLC nano HPLC (Thermo Fisher Scientific). Peptides were injected onto a 75 μm × 2 cm PepMap-C18 pre-column and resolved on a 75 μm × 50 cm RP-C18 EASY-Spray temperature controlled integrated column-emitter (Thermo Fisher Scientific). The mobile phases were: 2% ACN incorporating 0.1% FA (Solvent A) and 80% ACN incorporating 0.1% FA (Solvent B). A gradient from 5% to 35% solvent B was used over 4 h with a constant flow of 200 nL/min. The spray was initiated by applying 2.5 kV to the EASY-Spray emitter and the data were acquired on a Q-Exactive Orbitrap (Thermo Fisher Scientific) under the control of Xcalibur software in a data dependent mode selecting the 15 most intense ions for HCD-MS/MS. Raw MS data was processed using MaxQuant (version 1.3.0.5) (Cox & Mann, 2008).

### Mass Spectrometry Data Analysis

Putative readers were ranked based on the median of the intensity-based absolute quantification (iBAQ) values (Schwanhäusser *et al*, 2011). Briefly, for normalisation, iBAQ values were divided by the total sum of intensity of each sample and the median of normalised biological replicates was calculated. A value of 1×10^−9^ (a number 100 times smaller than the smallest value in the dataset) was then added to all median values to calculate fold changes.

### SDS-PAGE

Samples were prepared and validation pulldowns were performed as described in “Protein extraction” and “RNA pulldown”, respectively. Pulldown eluates and input controls were loaded in a 4-12% Bis-Tris gel and ran in NuPAGE MOPS SDS running buffer (Invitrogen) at 200V for 50 min in a XCell SureLock electrophoresis system (Thermo Fisher Scientific) connected to a Bio-Rad PowerPac (Bio-Rad).

### Western Blotting

Proteins resolved in SDS-PAGE were transferred onto a nitrocellulose membrane (Hybond ECL, Amersham) using NuPAGE Transfer Buffer (Invitrogen), for 2 h, at 250 mA, 4°C in a Mini Trans-Blot Cell (Bio-Rad). Membranes were blocked in 3% non-fat dry milk in TBS-T buffer for 30 min at room temperature. Membranes were incubated with primary antibody (mouse anti-FLAG M2, Sigma F1804) at 1:1000 dilutions in 3% milk-TBS-T overnight rotating at 4°C. Following three washes in TBST-T for 10 min each, membranes were incubated with secondary antibody diluted in milk/TBS-T for 1–2 hours at room temperature. Following 3 washes in TBS-T, bands were detected using Immobilon Western Chemiluminescent HRP Substrate (Millipore) according to manufacturer’s instructions. Medical X-ray films (Super Rx, Fuji) were exposed to luminescent membrane for the time required and the films were developed on a Compact X4 automatic X-ray film processor (Xograph Imaging Systems Ltd).

## ACKNOWLEDGEMENTS

We would like to thank the Gurdon Institute Media Kitchen for their support providing reagents and media. Tagged ALY-1 and ALY-2 strains were generated by the TransgeneOme project and provided by the *Caenorhabditis* Genetics Center – the latter is funded by NIH Office of Research Infrastructure Programs (P40 OD010440). This research was supported by a Wellcome Trust Senior Investigator award (104640/Z/14/Z) and Cancer Research UK award (C13474/A27826) to E.A.M; Conselho Nacional de Desenvolvimento Científico e Tecnológico doctorate scholarship (CNPq, Brazil - 205589/2014-6) to I.C.N; Wellcome Trust Senior Investigator award (209441/Z/17/Z) to S.B. The Miska laboratory is supported by core funding from the Wellcome Trust (092096/Z/10/Z, 203144/Z/16/Z) and Cancer Research UK (C6946/A24843). The Balasubramanian laboratory is supported by Cancer Research UK core (C9545/A19836) and programme award funding (C9681/A29214), and Herchel Smith Funds. To comply with Open Access requirements, the author has applied a CC BY public copyright licence to any Author Accepted Manuscript version arising from this submission.

## REFERENCES

Abbasi-Moheb L, Mertel S, Gonsior M, Nouri-Vahid L, Kahrizi K, Cirak S, Wieczorek D, Motazacker MM, Esmaeeli-Nieh S, Cremer K, et al (2012) Mutations in NSUN2 cause autosomal-Recessive intellectual disability. Am J Hum Genet 90: 847–855

Amos H & Korn M (1958) 5-Methyl cytosine in the RNA of Escherichia coli. BBA -Biochim Biophys Acta 29: 444–445

Angeles-Albores D, N. Lee RY, Chan J & Sternberg PW (2016) Tissue enrichment analysis for C. elegans genomics. BMC Bioinformatics 17: 366

Bensaddek D, Narayan V, Nicolas A, Brenes Murillo A, Gartner A, Kenyon CJ & Lamond AI (2016) Micro-proteomics with iterative data analysis: Proteome analysis in C. elegans at the single worm level. Proteomics 16: 381–392

Boccaletto P, Machnicka MA, Purta E, Pį Atkowski P-L, Zej Bagí Nski B-L, Wirecki TK, D. Crécy V, Crécy-Lagard C, Ross R, Limbach PA, et al (2017) MODOMICS: a database of RNA modification pathways. 2017 update. Nucleic Acids Res 46: 303–307

Brenner S (1974) The genetics of Caenorhabditis elegans. Genetics 77: 71–94

Chen X, Li A, Sun B-F, Yang Y, Han Y-N, Yuan X, Chen R-X, Wei W-S, Liu Y, Gao C-C, et al (2019) 5-methylcytosine promotes pathogenesis of bladder cancer through stabilizing mRNAs. Nat Cell Biol 21: 978–990

Cohn W.E. (1960) Pseudouridine, a carbon-carbon linked ribonucleoside in ribonucleic acids: isolation, structure, and chemical characteristics. J Biol Chem 235: 1488–1498

Cox J & Mann M (2008) MaxQuant enables high peptide identification rates, individualized p.p.b.-range mass accuracies and proteome-wide protein quantification. Nat Biotechnol 26: 1367–1372

Davis F & Allen FW (1957) Ribonucleic acids from yeast which contain a fifth nucleotide. J Biol Chem 227: 907–915

Dominissini D, Moshitch-Moshkovitz S, Schwartz S, Salmon-Divon M, Ungar L, Osenberg S, Cesarkas K, Jacob-Hirsch J, Amariglio N, Kupiec M, et al (2012) Topology of the human and mouse m6A RNA methylomes revealed by m6A-seq. Nature 485: 201–206

Freude K, Hoffmann K, Jensen L-R, Delatycki MB, Des Portes V, Moser B, Hamel B, Van Bokhoven H, Moraine C, Fryns J-P, et al (2004) Report Mutations in the FTSJ1 Gene Coding for a Novel S-Adenosylmethionine-Binding Protein Cause Nonsyndromic X-Linked Mental Retardation. Am J Hum Genet 75: 305–309

García-Vílchez R, Sevilla A & Blanco S (2019) Post-transcriptional regulation by cytosine-5 methylation of RNA. Biochim Biophys Acta - Gene Regul Mech: #pagerange#

Gatfield D & Izaurralde E (2002) REF1/Aly and the additional exon junction complex proteins are dispensable for nuclear mRNA export. J Cell Biol 159: 579–588

Guy MP, Shaw M, Weiner CL, Hobson L, Stark Z, Rose K, Kalscheuer VM, Gecz J & Phizicky EM (2015) Defects in tRNA Anticodon Loop 2′-O-Methylation Are Implicated in Nonsyndromic X-Linked Intellectual Disability due to Mutations in FTSJ1. Hum Mutat 36: 1176–1187

Van Haute L, Dietmann S, Kremer L, Hussain S, Pearce SF, Powell CA, Rorbach J, Lantaff R, Blanco S, Sauer S, et al (2016) Deficient methylation and formylation of mt-tRNAMet wobble cytosine in a patient carrying mutations in NSUN3. Nat Commun 7: 12039

Huber SM, van Delft P, Mendil L, Bachman M, Smollett K, Werner F, Miska E a. & Balasubramanian S (2015) Formation and Abundance of 5-Hydroxymethylcytosine in RNA. ChemBioChem 23: 752–755

Huber SM, Van Delft P, Tanpure A, Miska EA & Balasubramanian S (2017) 2′-O-methyl-5-hydroxymethylcytidine: A second oxidative derivative of 5-methylcytidine in RNA. J Am Chem Soc 139: 1766–1769

Kawarada L, Suzuki T, Ohira T, Hirata S, Miyauchi K & Suzuki T (2017) ALKBH1 is an RNA dioxygenase responsible for cytoplasmic and mitochondrial tRNA modifications. Nucleic Acids Res 45: 7401–7415

Khan MA, Rafiq MA, Noor A, Hussain S, Flores J V., Rupp V, Vincent AK, Malli R, Ali G, Khan FS, et al (2012) Mutation in NSUN2, which encodes an RNA methyltransferase, causes autosomal-recessive intellectual disability. Am J Hum Genet 90: 856–863

Komara M, Al-Shamsi AM, Ben-Salem S, Ali BR & Al-Gazali L (2015) A Novel Single-Nucleotide Deletion (c.1020delA) in NSUN2 Causes Intellectual Disability in an Emirati Child. J Mol Neurosci 57: 393–399

Kuersten S, Segal SP, Verheyden J, LaMartina SM & Goodwin EB (2004) NXF-2, REF-1, and REF-2 Affect the Choice of Nuclear Export Pathway for tra-2 mRNA in C. elegans. Mol Cell 14: 599–610

Longman D, Johnstone IL & Cáceres JF (2003) The Ref / Aly proteins are dispensable for mRNA export and development in Caenorhabditis elegans The Ref / Aly proteins are dispensable for mRNA export and development in Caenorhabditis elegans. RNA 9: 881–891

Maris C, Dominguez C & Allain FHT (2005) The RNA recognition motif, a plastic RNA-binding platform to regulate post-transcriptional gene expression. FEBS J 272: 2118–2131

Navarro IC, Tuorto F, Jordan D, Legrand C, Price J, Braukmann F, Hendrick AG, Akay A, Kotter A, Helm M, et al (2020) Translational adaptation to heat stress is mediated by RNA 5-methylcytosine in Caenorhabditis elegans. EMBO J: 1–18

Païs de Barros J-P, Keith G, El Adlouni C, Glasser A-L, Mack G, Dirheimer G & Desgrès J (1996) 2′-O-methyl-5-formylcytidine (f5Cm), a new modified nucleotide at the ‘wobble’ position of two cytoplasmic tRNAs Leu (NAA) from bovine liver. Nucleic Acids Res 24: 1489–1496

Ramser J, Winnepenninckx B, Lenski C, Errijgers V, Platzer M, Schwartz CE & Meindl A (2004) A splice site mutation in the methyltransferase gene FTSJ1 in Xp11.23 is associated with non-syndromic mental retardation in a large Belgian family (MRX9). J Med Genet 41: 679–683

Sarov M, Murray JI, Schanze K, Pozniakovski A, Niu W, Angermann K, Hasse S, Rupprecht M, Vinis E, Tinney M, et al (2012) A genome-scale resource for in vivo tag-based protein function exploration in C. elegans. Cell 150: 855–866

Schwanhäusser B, Busse D, Li N, Dittmar G, Schuchhardt J, Wolf J, Chen W & Selbach M (2011) Global quantification of mammalian gene expression control. Nature 473: 337–342

Shi H, Wei J & He C (2019) Where, When, and How: Context-Dependent Functions of RNA Methylation Writers, Readers, and Erasers. Mol Cell 74: 640–650

Takano K, Nakagawa E, Inoue K, Kamada F, Kure S, Goto YI, Inazawa J, Kato M, Kubota T, Kurosawa K, et al (2008) A loss-of-function mutation in the FTSJ1 gene causes nonsyndromic x-linked mental retardation in a Japanese family. Am J Med Genet Part B Neuropsychiatr Genet 147: 479–484

Takemoto C, Spremulli LL, Benkowski LA, Ueda T, Yokogawa T & Watanabe K (2009) Unconventional decoding of the AUA codon as methionine by mitochondrial tRNA Met with the anticodon f 5 CAU as revealed with a mitochondrial in vitro translation system. Nucleic Acids Res 37: 1616–1627

Tanpure AA & Balasubramanian S (2017) Synthesis and Multiple Incorporations of 2′-O-Methyl-5-hydroxymethylcytidine, 5-Hydroxymethylcytidine and 5-Formylcytidine Monomers into RNA Oligonucleotides. ChemBioChem 18: 2236–2241

Yang X, Yang Y, Sun B-F, Chen Y-S, Xu J-W, Lai W-Y, Li A, Wang X, Bhattarai DP, Xiao W, et al (2017) 5-methylcytosine promotes mRNA export — NSUN2 as the methyltransferase and ALYREF as an m5C reader. Cell Res: 1–20

Yang Y, Wang L, Han X, Yang WL, Zhang M, Ma HL, Sun BF, Li A, Xia J, Chen J, et al (2019) RNA 5-Methylcytosine Facilitates the Maternal-to-Zygotic Transition by Preventing Maternal mRNA Decay. Mol Cell 75: 1188–1202.e11

